# Exploring the Dose-Dependency of After-Effects: A Computational Model for Theta-Burst Transcranial Magnetic Stimulation

**DOI:** 10.1101/2023.07.03.547540

**Authors:** Ke Ma, Sung Wook Chung, Stephan M. Goetz

## Abstract

Transcranial magnetic stimulation (TMS) is a non-invasive neurostimulation and neuromodulation technique that is widely applied in brain research and clinical applications. However, the optimal parameters of neuromodulating TMS protocols describing the specific rhythms, such as number of pulses, frequency, and stimulation strength, are widely unknown. Improving previous rather limited and ad-hoc models, we aimed to investigate the dose-dependency of theta-burst stimulation (TBS) protocols with a more elaborate but still parsimonious quantitative model representing the non-linearities of the mechanisms of synaptic plasticity and metaplasticity during repetitive magnetic stimulation. Our model, which considers the interaction between facilitatory and inhibitory processes, successfully reproduced results from TBS experiments and provide testable predictions for prolonged TBS protocols. Moreover, we suggested that the activation of kinases and phosphatases could be potential candidates for later TMS modelling research. Although this model still simplifies the complex dynamics of cellular and molecular processes, it offers a starting basis for future studies to incorporate more intricate mechanisms. By further refining our understanding of the underlying mechanisms and improving the accuracy of prediction models, we can advance the efficacy and clinical application of TBS protocols in various neurological and psychiatric conditions.

## 1 Introduction

Transcranial magnetic stimulation (TMS) is a non-invasive brain stimulation technique using brief strong electromagnetic pulses to write signals into neural circuits, most prominently of the human cortex (Barker et al. 1985). Certain rhythms can further modulate, i.e., influence how a circuit processes endogenous signals (Di Lazzaro et al. 2012; Di Lazzaro et al. 2018). It finds extensive applications in both experimental brain research and clinical treatment, including the therapy of major depression, obsessive-compulsive disorder, addiction, migraine, stroke, and epilepsy as well as in diagnostic procedures (Beynel et al. 2019; Chung et al. 2015; Eldaief et al. 2013; Luan et al. 2014; Mantovani et al. 2010; Rossini et al. 2015; Valero-Cabré et al. 2017; Wagner et al. 2007). The improvement of device technology allows more intricate accelerated neuromodulation protocols (Goetz and Deng 2017). Theta-burst stimulation (TBS), for example, is a highly effective and quick neuromodulatory protocol for TMS proposed by Huang et al. (2005). The primary motor cortex often serves as a model circuit in many quantitative studies, where motor-evoked potentials (MEP) in response to constant single-pulse TMS can reveal TBS-induced excitability changes.

Since the introduction of TBS, most studies have used either intermittent thetaburst stimulation with 600 pulses (iTBS600) or continuous theta-burst stimulation with 600 pulses (cTBS). These protocols commonly employ a pulse strength of 80 % of the active motor threshold, along with a specific pattern consisting of three pulses at 50 Hz per burst and a certain number of bursts at 5 Hz per train (Chung et al. 2016; Corp et al. 2020; Turi et al. 2021). It has been observed that iTBS600 generally leads to an increase in MEPs, while cTBS600 tends to decrease MEPs (Chung et al. 2016; Suppa et al. 2016). Furthermore, a few studies explored variations in the number of pulses within TBS protocols and demonstrated that prolonged TBS protocols exceeding 600 pulses can reverse the effect of the standard TBS protocols, a phenomenon termed dose-dependency (Gamboa et al. 2010; McCalley et al. 2021; Moisset et al. 2015). Huang et al. (2011) first developed a theoretical model based on the hypothesis of synaptic plasticity. The model successfully reproduced the after-effects of cTBS600 and iTBS600 and suggested that the calcium dynamics at post-synapses play a bi-directional regulatory role in these after-effects. Recent work fixed a few discrepancies and re-calibrated the model (Ma et al. 2022).

Several studies have suggested that homeostatic metaplasticity could potentially explain the reversal of after-effects induced by TMS, as it has the ability to modulate synaptic plasticity in a higher-order manner in case of instability (Abraham 2008; Gamboa et al. 2011; Hamada et al. 2008; Huang et al. 2010; Murakami et al. 2012; Todd et al. 2009). However, the available rather simple TBS models fail to reproduce the after-effects induced by TBS protocols involving more than 600 pulses (Huang et al. 2011; Ma et al. 2022).

To address this limitation, Fung and Robinson (2014) developed a TBS model that incorporates a calcium-dependent learning function derived from Shouval et al. (2002) within a single-population neural field model constructed by Robinson (2011). This model introduces metaplasticity by involving varying calcium conductance for N-methyl-D-aspartate receptors (NMDAR) proposed by Yeung et al. (2004). Although this model can simulate the after-effects of cTBS and iTBS with any given number of pulses, it exhibits inconsistencies with several experimental results and requires further evaluation (Wilson et al. 2018). Subsequently, Wilson et al. (2016) extended the single-population neural field model to a triple-population neural field model and explored different patterns of iTBS and cTBS using only 600 pulses. Similarly, an improved model developed by Wilson et al. (2021) only simulates the standard cTBS600 and iTBS600. Despite including biological mechanisms and phenomenological observations, these models share a significant limitation in that none of them can reproduce the existing neuromodulatory after-effects induced by TBS with different numbers of pulses. Accordingly, there is a pressing need for a quantitative model that can provide potential insights into the underlying mechanisms behind the diverse neuromodulatory after-effects induced by different TBS protocols, reproduce experimental findings, and offer reasonable predictions to guide further TBS research and protocol enhancement.

This study aims to develop a model that specifically focuses on the dose-dependent after-effects of TBS when applied to the primary motor cortex. The model will provide predictions for untested protocols, offering valuable experimental guidance. The following text will present a detailed procedure for constructing the model, along with the corresponding assumptions used. Furthermore, simulation results will be compared with experimental findings, enabling a comprehensive discussion of the underlying mechanisms. Finally, this article will conclude with a summary of the overall reproducibility and predictability of the proposed model.

## 2 Mathematical model

TBS predominantly modulates I-waves rather than D-waves in the epidural recording of descending volleys evoked by single-pulse TMS, indicating that the induced neuromodulation may modify the synaptic transmission between neurons of cortical circuits (Di Lazzaro et al. 2008, 2005). Di Lazzaro et al. (2010) and Huang et al. (2017) suggested that the after-effects of TMS are mediated by cellular physiological mechanisms, particularly calcium-dependent bi-directional synaptic plasticity. Moreover, the activation of NMDARs, which are major ionotropic glutamate receptors permeable to calcium ions, can significantly influence the after-effects induced by TBS (Huang et al. 2007; Liepert et al. 1997). Therefore, we apply the mechanisms of calcium-dependent plasticity at glutamatergic synapses to model the after-effect of TBS. The overall model has four subsystems, describing the dynamics of calcium concentration, calcium-induced signalling cascades, changes of MEP learning, and metaplasticity induction.

### Intracellular calcium dynamics

Huang et al. (2007) demonstrated that the after-effects of TBS depend on the activation of NMDARs. Hence, in our model, we focus on the calcium influx through NMDARs as the primary source of calcium. TMS is known to stimulate various types of intracortical neurons in the superficial regions of the cerebral cortex. As a result, both presynaptic and postsynaptic neurons within a circuit are stimulated in the target region. The elevation of the postsynaptic membrane potential leads to the opening of NMDARs, while glutamate is released pre-synaptically (Mayer et al. 1984). For simplicity, we assume that the calcium influx is instantaneously activated by TMS, resulting in calcium transients represented by [Ca^2+^]_i_, and each stimulus is denoted by an impulse input to the system. The postsynaptic calcium concentration can be approximately expressed as

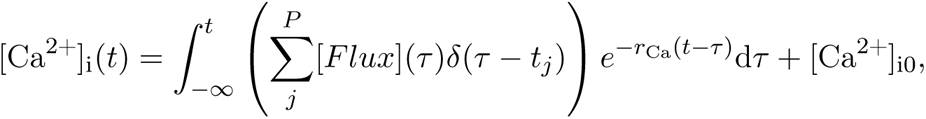

where *δ*(*t*) denotes the Dirac delta distribution, *t_j_*is the time of the *j*-th pulse administered, *r*_Ca_ represents the decay rate of intracellular calcium concentration, [Ca^2+^]_i0_ is the resting calcium concentration, [*Flux*](*t*) refers to the varying rate of calcium influx at time *t*, and *P* is the total number of pulses applied. By solving this integral equation, the dynamics of intracellular calcium concentration follows

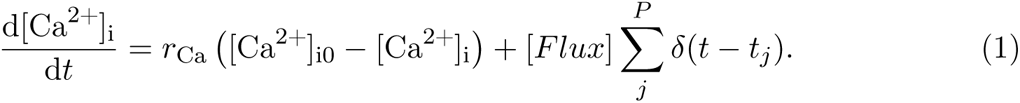

As suggested by Shouval et al. (2002), the decay constant of intracellular calcium is 50 ms. Accordingly, the calcium decay rate in this study follows *r*_Ca_ = 20 s*^−^*^1^. In addition, the resting calcium concentration is 100 nM (Berridge et al. 2000).

### Facilitation and inhibition substances

Several studies have provided converging evidence to support the idea that the long-term potentiation (LTP) and long-term depression (LTD) of synaptic plasticity among glutamatergic neurons are heavily reliant on postsynaptic calcium influx mediated by NMDARs (Artola et al. 1990; Bear and Malenka 1994; Bi and Poo 1998; Kirkwood et al. 1993). Calcium, as a second messenger, triggers downstream calcium-sensitive protein kinases and phosphatases in the postsynaptic neurons, bi-directionally regulating synaptic plasticity (Berridge 1998). Two prominent hypotheses have been proposed to explain the role of calcium in synaptic plasticity. First, the calcium amplitude hypothesis suggests that the postsynaptic calcium level determines the direction of induced plasticity (Artola and Singer 1993). Specifically, a higher calcium concentration can trigger long-term potentiation, while a moderate calcium concentration is necessary for long-term depression (Artola et al. 1990). Secondly, the calcium temporal patterns hypothesis emphasises the importance of the time course of calcium elevation. For instance, Yang et al. (1999) demonstrated that a slower, lower calcium elevation consistently induces long-term depression, whereas a faster, higher calcium transient results in long-term potentiation. Moreover, Evans and Blackwell (2015) suggested that the calcium pattern hypothesis does not contradict the calcium amplitude hypothesis, but rather it focuses more on the temporal patterns of calcium transients.

Therefore, this model uses two substances, termed [*Faci*] and [*Inhi*], to represent the postsynaptic facilitatory and inhibitory signalling cascades, respectively. Taking into account the temporal patterns and the existence of two thresholds for calcium signalling observed in experiments, this study uses a soft dose-response activation function, known as the Hill function, to describe the relationships between [Ca^2+^]_i_ and the substances. The inhibition substance overall exhibits higher sensitivity at low calcium concentrations, while the facilitation substance shows higher sensitivity at high calcium concentrations. We assume that the production rates of these substances solely depend on [Ca^2+^]_i_. Thus, their dynamics can be written as

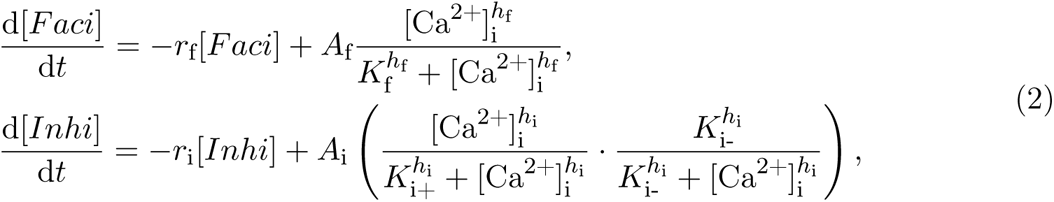

where *r*_f_ and *r*_i_ are the decay rates of substances, *A*_f_ and *A*_i_ represent the maximum production rates, *h*_f_ and *h*_i_ are the Hill coefficients, *K*_f_ is the half-maximal values of [Ca^2+^]_i_ for facilitation, and *K*_i+_ and *K*_i-_ are the half-maximal values of [Ca^2+^]_i_ for ascending and descending phases of inhibition substances respectively. Both substances are in arbitrary units.

### MEP learning procedure

*M*_net_ is the maximum/minimum value of the normalised MEP measured after the intervention in this study. The facilitation substance increases the amplitude of MEP, while the inhibition substance decreases it. Both of them compete during the intervention. We base the MEP learning procedure on the Michaelis–Menten enzyme kinetics, which follows

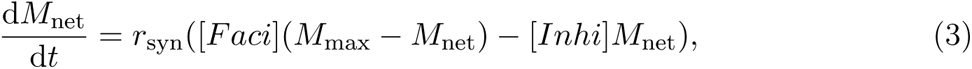

where *r*_syn_ is the learning rate of MEP and *M*_max_ the upper limit for the range of *M*_net_. The initial value of *M*_net_ is one given in arbitrary units, which could be regarded as the baseline MEP before the intervention, and we set *M*_max_ = 2 here.

### Calcium influx rate and metaplasticity

Furthermore, several lines of evidence indicate that the TMS-induced after-effects depend on the prior history of cortical activity, suggesting the involvement of metaplasticity during TMS (Gamboa et al. 2011, 2010; Hamada et al. 2008; Huang et al. 2010; Murakami et al. 2012; Todd et al. 2009). Studies on tetanic synapses stimulation have demonstrated that metaplasticity is dependent on the historical activation of NMDARs and synaptic changes (Huang et al. 1992; Ngezahayo et al. 2000; Tokay et al. 2014). For the sake of simplicity, this study does not incorporate the dynamics of conductance of NMDARs as modelled by Yeung et al. (2004). Instead, we represent the changes in NMDARs by varying the rates of calcium influx. Therefore, the calcium influx rate follows

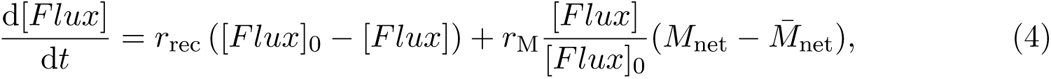

where *r*_rec_ is the recovery rate of calcium influx rate, [*Flux*]_0_ denotes the naive calcium influx rate and 1*/*[*Flux*]_0_ acts as a scaling constant, and *r*_M_ is the induction rate of metaplasticity on [*Flux*]. We set [*Flux*]_0_ = 20 µM*/*s to let the intracellular calcium concentration vary in a reasonable range (Berridge et al. 2000; Fung and Robinson 2013; Shouval et al. 2002). 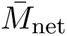 is the time average of *M*_net_ as the historical memory of the system, which is given by

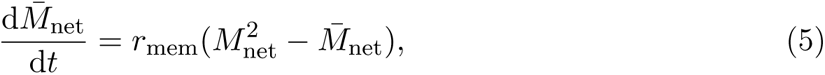

where *r*_mem_ is the rate of memory establishment for metaplasticity. This equation follows the sliding threshold equation proposed by Law and Cooper (1994).

### After-effect curve equation

The net changes of normalised MEPs can be calculated as Δ*M*_net_ = *M*_net_ *−* 1. It has been demonstrated that MEPs will persist for a certain duration before returning to their baseline values (Di Lazzaro et al. 2002a; Huang et al. 2005). We propose a three-phase after-effect curve equation, consisting of rising, preservation, and decay phases. The after-effect curve follows

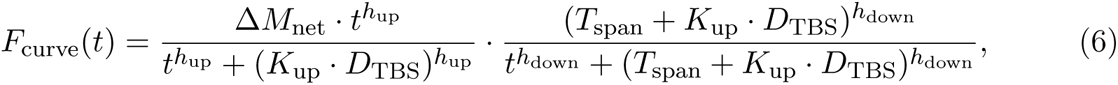

where *h*_up_ and *h*_down_ are power coefficients for rising and decay phases respectively, *K*_up_ is the ratio coefficient for the rising phase, *D*_TBS_ denotes the stimulation duration of a TBS session. According to the study of Huang et al. (2011), the time span of the preservation phase, *T*_span_, should depend on the number of pulses applied and follow

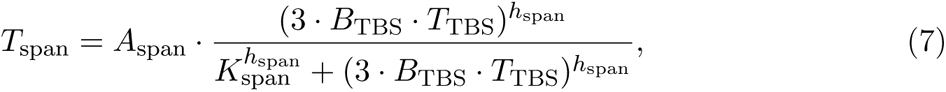

where *A*_span_ represents the maximum time span of the preservation phase, *K*_span_ is the half-maximal value of time, and *h*_span_ is the power coefficient. *B*_TBS_ denotes the number of bursts per train, and *T*_TBS_ is the number of trains per session.

## 3 Methodology for parameter calibration

We collected the TBS experimental data, specifically the reported mean values of MEPs, from studies listed in the supplementary document (see Supplementary Material). Where data were not reported numerically, we extracted them through plot digitisation (Rohatgi 2017). All of these studies used biphasic pulses and set the stimulus strength to 80% active motor threshold. For iTBS, the number of administered pulses were 600, 1200, and 1800, while cTBS used pulse numbers of 300, 600, 1200, and 1800. Before calibration, all measurements of the TBS-induced after-effects on MEPs were normalised by the baseline MEP (specifically the MEP measured before TBS intervention). Furthermore, we evaluated the delta MEP by subtracting 1 in this database.

The proposed model requires calibration of a set of parameters denoted as **X**. Suppose that there are *Q* measurement patches for *Q* TBS protocols, (i.e., each data patch includes MEP measurements of the same TBS protocol), the *i*-th TBS patch is associated with a cost function *J_i_*(**X**) for *i*-th TBS protocol. The overall cost function is obtained by summing these individual cost functions together, yielding a scalar value. The complete cost function follows

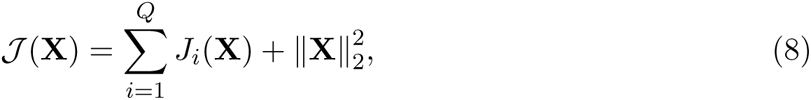

where 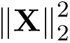 represents the squared p2 norm of the parameter vector **X**, serving as a regularisation term. The cost function *J_i_*(**X**) for the *i*-th TBS patch is given by

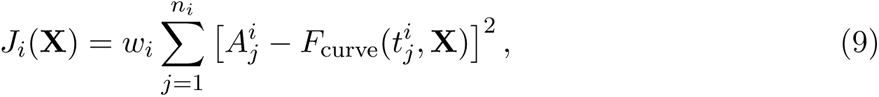

where 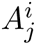 is the *j*-th measurement in the *i*-th TBS patch measured at time 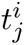. In addition, because of the imbalanced data size for each patch, we introduce 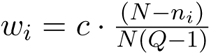 to weigh the cost of each patch, where *N* is the total number of measurements in the database, *c* is an arbitrary scaling constant, and *n_i_* is the number of measurements in the *i*-th patch such that 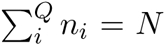. Hence, the sum of these weights equals *c*. The cost function quantifies the discrepancy between the experimental measurements and the model prediction obtained using the function 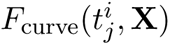. The goal of the calibration process is to minimise this cost function in order to find the optimal values for the parameter vector **X**.

The calibration of the proposed model parameters is subject to certain constraints, specifically, all parameters should be real positive numbers. We formulate a non-convex and nonlinear optimisation problem for optimally calibrating the parameters as

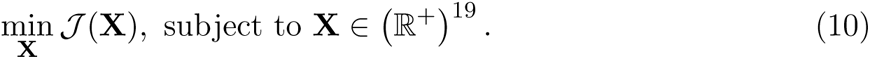

We used the global particle swarm optimisation algorithm implemented in Matlab (v2022b, The Mathworks, USA) with 500 workers in the swarm for solving Equations 8 and 10 to avoid the problem of poor convergence and local minima with *c* = 10. Due to the nonlinearity of this cost function, global optimality is not necessarily guaranteed. The calibrated parameters are listed in Table 1. This study uses the 4-th order Runge–Kutta method with a time step of 0.01 s to calculate the dynamics of the system. The model codes are provided in the supplementary material (see Supplementary Material).

**Table 1:**
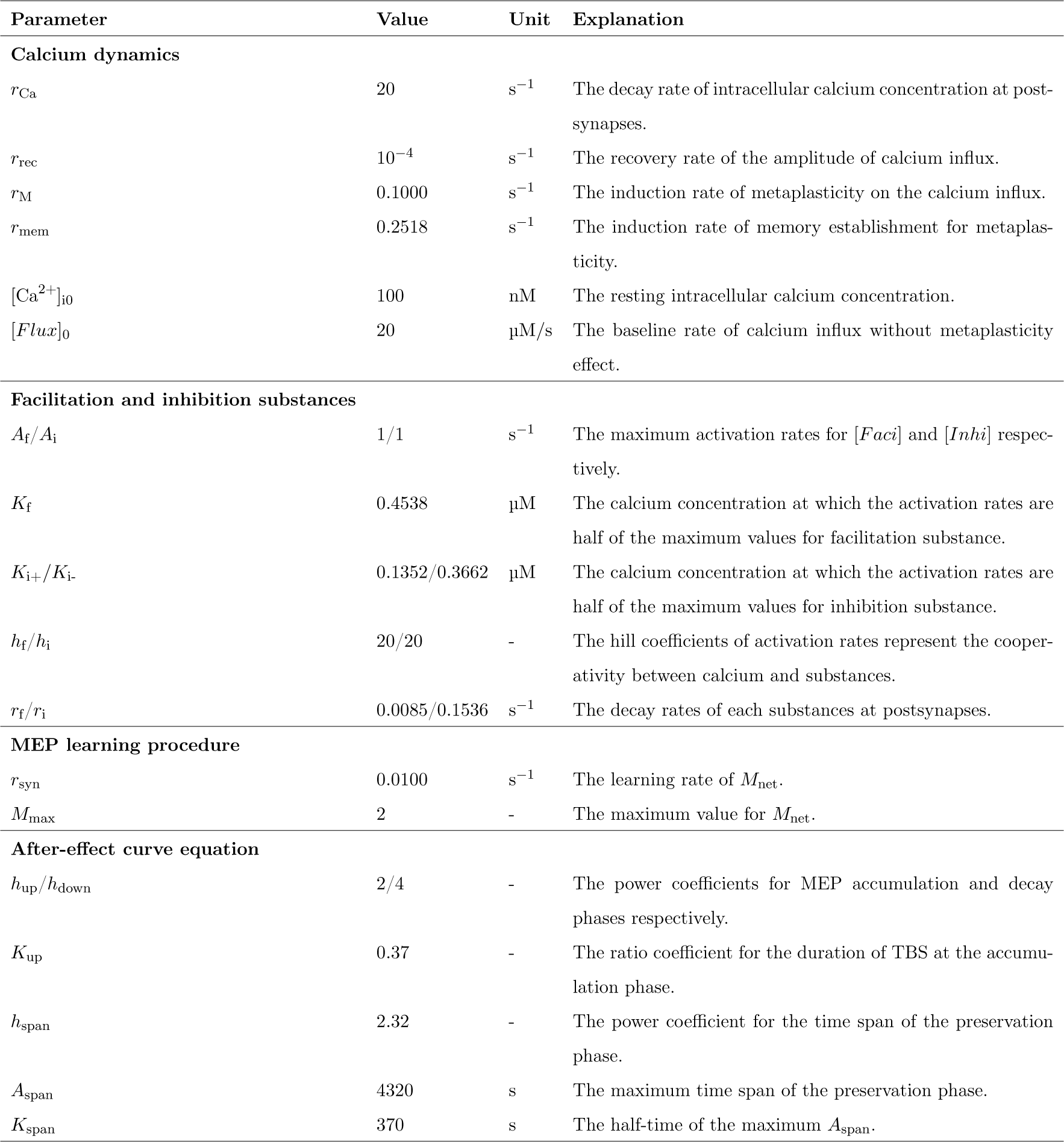
Summary of the calibrated parameters used in the nonlinear system.

| Parameter | Value | Unit | Explanation |
| --- | --- | --- | --- |
| **Calcium dynamics** |  |  |  |
| $r_{Ca}$ | 20 | $s^{-1}$ | The decay rate of intracellular calcium concentration at post-synapses. |
| $r_{rec}$ | $10^{-4}$ | $s^{-1}$ | The recovery rate of the amplitude of calcium influx. |
| $r_M$ | 0.1000 | $s^{-1}$ | The induction rate of metaplasticity on the calcium influx. |
| $r_{mem}$ | 0.2518 | $s^{-1}$ | The induction rate of memory establishment for metaplasticity. |
| $[Ca^{2+}]_{i0}$ | 100 | nM | The resting intracellular calcium concentration. |
| $[Flux]_0$ | 20 | $\mu M/s$ | The baseline rate of calcium influx without metaplasticity effect. |
| **Facilitation and inhibition substances** |  |  |  |
| $A_f/A_i$ | 1/1 | $s^{-1}$ | The maximum activation rates for $[Faci]$ and $[Inhi]$ respectively. |
| $K_f$ | 0.4538 | $\mu M$ | The calcium concentration at which the activation rates are half of the maximum values for facilitation substance. |
| $K_{i+}/K_{i-}$ | 0.1352/0.3662 | $\mu M$ | The calcium concentration at which the activation rates are half of the maximum values for inhibition substance. |
| $h_f/h_i$ | 20/20 | - | The hill coefficients of activation rates represent the cooperativity between calcium and substances. |
| $r_f/r_i$ | 0.0085/0.1536 | $s^{-1}$ | The decay rates of each substances at postsynapses. |
| **MEP learning procedure** |  |  |  |
| $r_{syn}$ | 0.0100 | $s^{-1}$ | The learning rate of $M_{net}$. |
| $M_{max}$ | 2 | - | The maximum value for $M_{net}$. |
| **After-effect curve equation** |  |  |  |
| $h_{up}/h_{down}$ | 2/4 | - | The power coefficients for MEP accumulation and decay phases respectively. |
| $K_{up}$ | 0.37 | - | The ratio coefficient for the duration of TBS at the accumulation phase. |
| $h_{span}$ | 2.32 | - | The power coefficient for the time span of the preservation phase. |
| $A_{span}$ | 4320 | s | The maximum time span of the preservation phase. |
| $K_{span}$ | 370 | s | The half-time of the maximum $A_{span}$. |

## 4 Results

### Substance growth rates

Equation 2 provides a description of the substance dynamics. The growth rates for [*Faci*] and [*Inhi*] are only determined by [Ca^2+^]i and are modelled by the Hill equations, as illustrated in Figure 1. Notably, the inhibition substance has a higher growth rate at low [Ca^2+^]i, while the facilitation substance has a higher growth rate at high [Ca^2+^]i. The cross-over point between them occurs approximately at [Ca^2+^]i = 0.41 µM. Upon examining the calibrated parameters presented in Table 1s, it is noteworthy that the decay rate of the facilitation substance (*r*_f_ = 0.0085 s*^−^*^1^) is significantly lower than that of the inhibition substance (*r*_i_ = 0.1536 s*^−^*^1^). This implies that the inhibition substances immediately respond to changes in [Ca^2+^]_i_, while the facilitation substance accumulates gradually as [Ca^2+^]_i_ increases over time.

**Figure 1:**
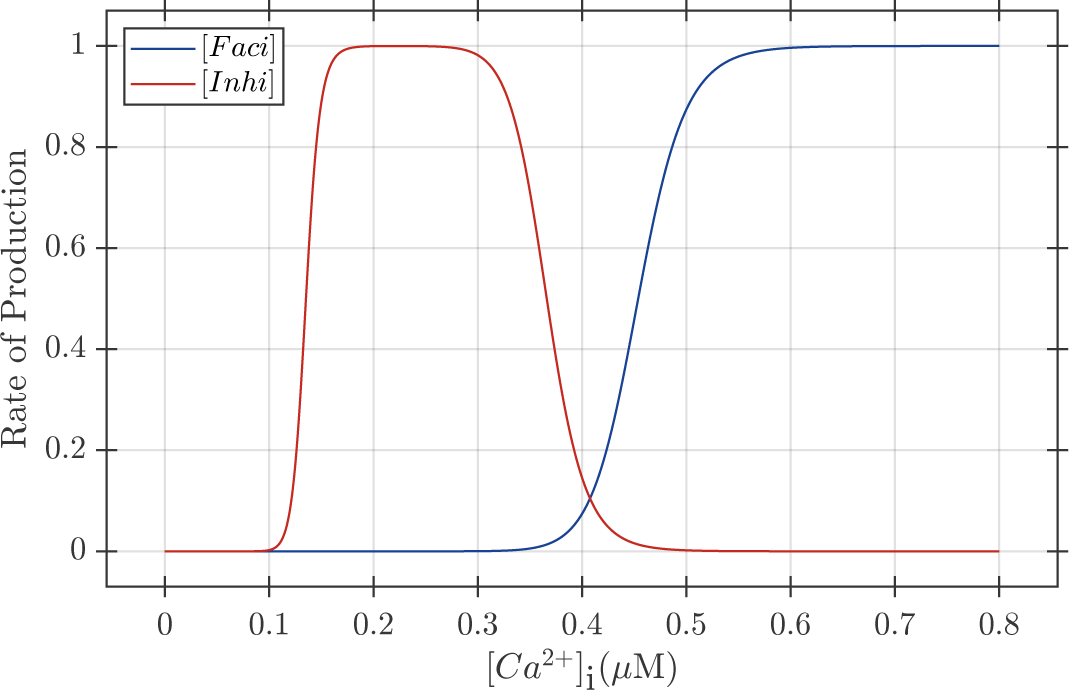
Growth rates of facilitation and inhibition substances depending on the intracellular calcium concentration [Ca^2+^]_i_.

### Simulation results for iTBS

Figure 2 shows the simulation results for iTBS with different numbers of pulses. Figures 2 (**A**) to (**D**) depict the system responses to a continuous iTBS application. Figure 2 (**D**) illustrates the net MEP effects for iTBS600, iTBS1200, and iTBS1800. As Figure 2 (**C**) demonstrates, fluctuations in [*Inhi*] occur in response to changes in [Ca^2+^]i, while [*Faci*] gradually accumulates in response to the elevation of [Ca^2+^]i.

**Figure 2:**
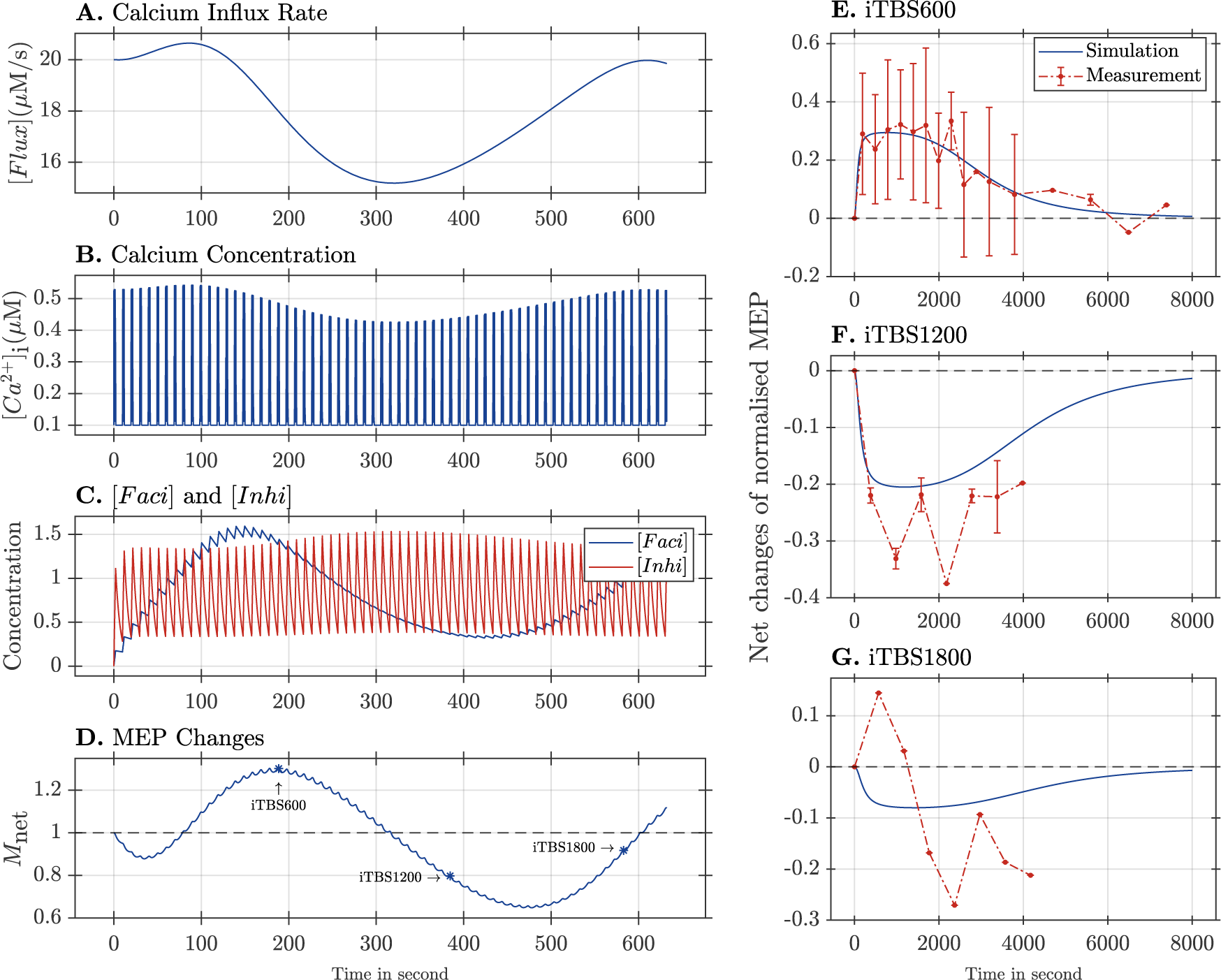
Simulation results for iTBS with 600, 1200, and 1800 pulses. (**A**) shows the calcium influx rate changes in response to the stimulus; (**B**) shows intracellular calcium concentration dynamics during the stimulation period; (**C**) shows the dynamics of facilitation and inhibition substances; (**D**) shows the net changes in MEP after stimulation; (**E**) shows the after-effect curve after iTBS600; (**F**) shows the after-effect curve after iTBS1200; and (**G**) shows the after-effect curve after iTBS1800. In panels (**E**) to (**G**), red dots represent the mean value of MEPs measured at specific time points, with red bars indicating the corresponding standard deviations, and the blue curves represent simulated after-effect curves.

At the onset of the iTBS protocol, there is an increase in the rate of calcium influx, leading to a rise in the peak amplitude of [Ca^2+^]_i_. Due to the higher sensitivity of the inhibition substance compared to the facilitation substance at low calcium concentrations, [*Inhi*] initially surpasses [*Faci*] and then introduces an inhibitory effect (*M* net *<* 1) to *M* net as observed in Figure 2 (**D**). Additionally, this also causes an inhibitory change in the historical memory system, 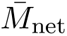, which increases the rate of calcium influx per stimuli for the following intervention. As [Ca^2+^]_i_ increases, the production rate of [*Faci*] increases, leading to a fast accumulation over time and a facilitatory effect on *M*_net_ afterwards. However, as [Ca^2+^]_i_ decreases and approaches the cross-over concentration because of the historical facilitation, the production rate of [*Faci*] decreases, leading to an inability to sustain its level. Meanwhile, [*Inhi*] can still respond to changes in [Ca^2+^]_i_. Consequently, an inhibitory effect is induced on *M*_net_. Therefore, the historical memory system over the past period can regulate the calcium influx responses to upcoming stimuli, which provide a feedback mechanism to control and restrict [Ca^2+^]_i_ levels and the induced effect on *M*_net_.

Figure 2 (**D**) shows that iTBS600 leads to a facilitated after-effect on MEP (Figure 2 (**E**)). On the other hand, iTBS1200 and iTBS1800 induce inhibited after-effects on MEP, as shown in Figures 2 (**F**) and (**G**). These findings align with previous studies conducted by Gamboa et al. (2010) and McCalley et al. (2021). Although the peak amplitude of the simulation curve for iTBS1800 does not precisely match the experimental results, the simulation results for all iTBS protocols successfully capture the polarities and durations observed in the experiments.

### Simulation results for cTBS

In Figure 3, the stimulation results for cTBS with different numbers of pulses applied are presented. Figures 3 (**A**) to (**D**) illustrate the system responses to a continuous cTBS application. In comparison to iTBS, Figure 3 (**B**) demonstrates that the dynamics of [Ca^2+^]_i_ exhibit smoother behaviour due to the continuous stimulus application without inter-train intervals characteristic of iTBS protocols. Moreover, the varying range of [Ca^2+^]_i_ to cTBS is also smaller than that to iTBS. Similar to iTBS, the initial stimuli of cTBS induce the inhibition substance first, followed by a gradual increase in the facilitation substance.

**Figure 3:**
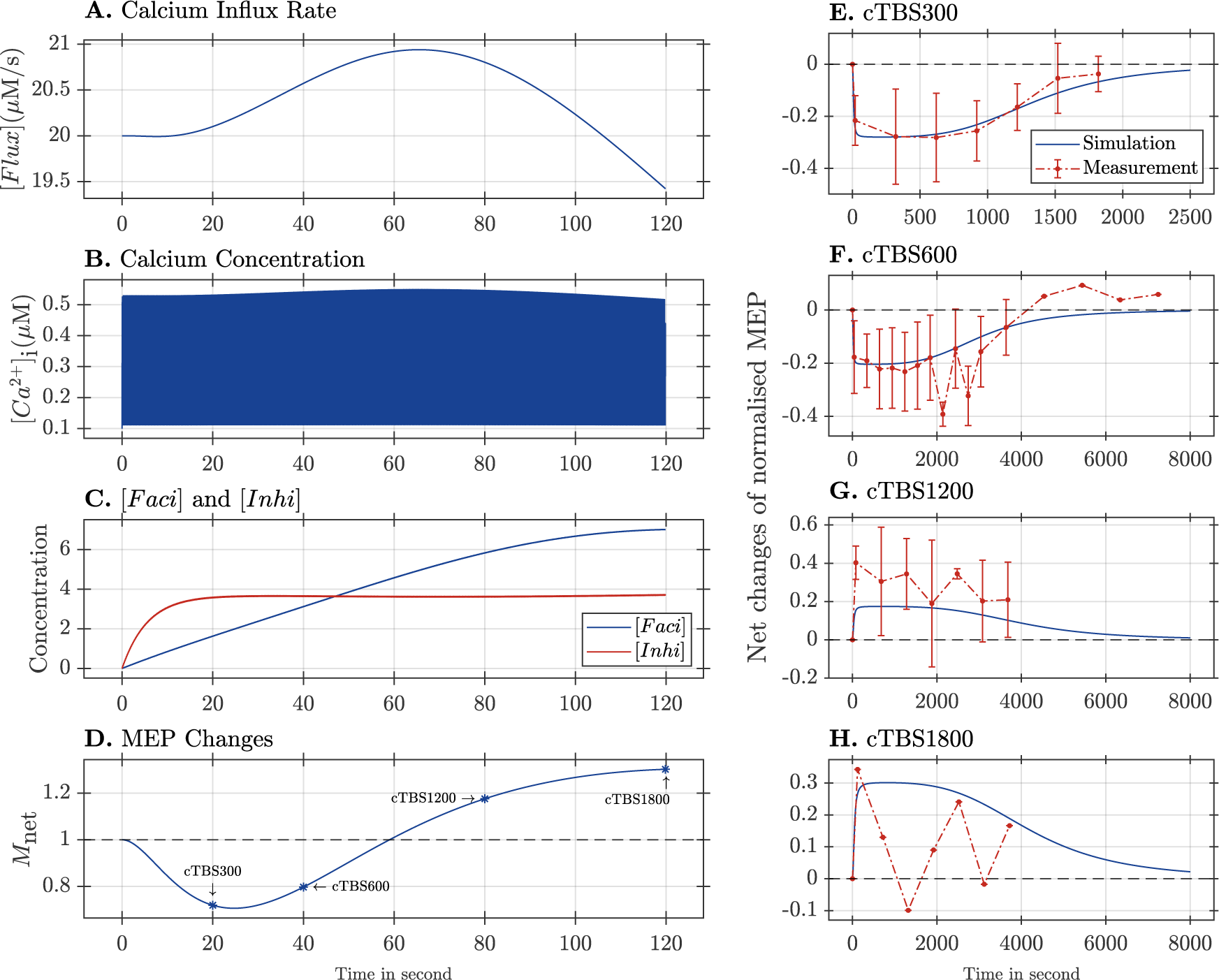
Simulation results for cTBS with 300, 600, 1200, and 1800 pulses. (**A**) shows the calcium influx rate changes in response to the stimulus; (**B**) shows intracellular calcium concentration dynamics during the stimulation period; (**C**) shows the dynamics of facilitation and inhibition substances; (**D**) shows the net changes of MEP after stimulation; (**E**) shows the after-effect curve after cTBS300; (**F**) shows the after-effect curve after cTBS600; (**G**) shows the after-effect curve after cTBS1200; (**H**) shows the after-effect curve after cTBS1800. In panels (**E**) to (**H**), red dots represent the mean value of MEPs measured at specific time points, with red error bars indicating the corresponding standard deviations, and the blue curves represent simulated after-effect curves.

Because cTBS has a denser distribution of pulses, this allows [Ca^2+^]_i_ to maintain a level slightly higher than the cross-over point. The slow decay of [*Faci*] allows for its accumulation when its growth rate is not significant during periods of relatively low [Ca^2+^]_i_, eventually surpassing [*Inhi*]. Therefore, due to the slow accumulation of [*Faci*], both cTBS300 and cTBS600 inhibit and decrease the post-intervention MEPs in response to probing pulses. However, prolonged inhibition of *M*_net_ will promote the calcium influx rate, increase [Ca^2+^]_i_, and induce a facilitatory effect on MEPs. For example, the polarity of the after-effects is reversed by cTBS1200 and cTBS1800 (Figure 3 (**D**)). Moreover, prolonged facilitation of *M*_net_ will once again suppress the calcium influx rate, decrease [Ca^2+^]_i_, and induce an inhibitory effect on MEP. Figures 3 (**E**) to (**H**) illustrate the simulated after-effects induced by cTBS protocols. These simulation results successfully capture the experienced changes in MEPs following stimulation, including the polarities, amplitudes, and durations.

### Simulation results for prediction

Figure 4 presents the prediction results obtained from the continuous application of cTBS and iTBS protocols. The system’s response to cTBS dampens faster and has fewer oscillations before going zero compared to iTBS. This discrepancy might be attributed to the structural differences between these two protocols. Particularly, cTBS employs a denser pulse array, delivering more pulses to the target brain area within the same period as iTBS. This denser pulse delivery of cTBS may explain the rapid damping of MEP changes in responses to this protocol. However, it is important to note that the prediction results should be interpreted with caution, as this nonlinear system was only fitted to a limited database comprising several TBS protocols.

**Figure 4:**
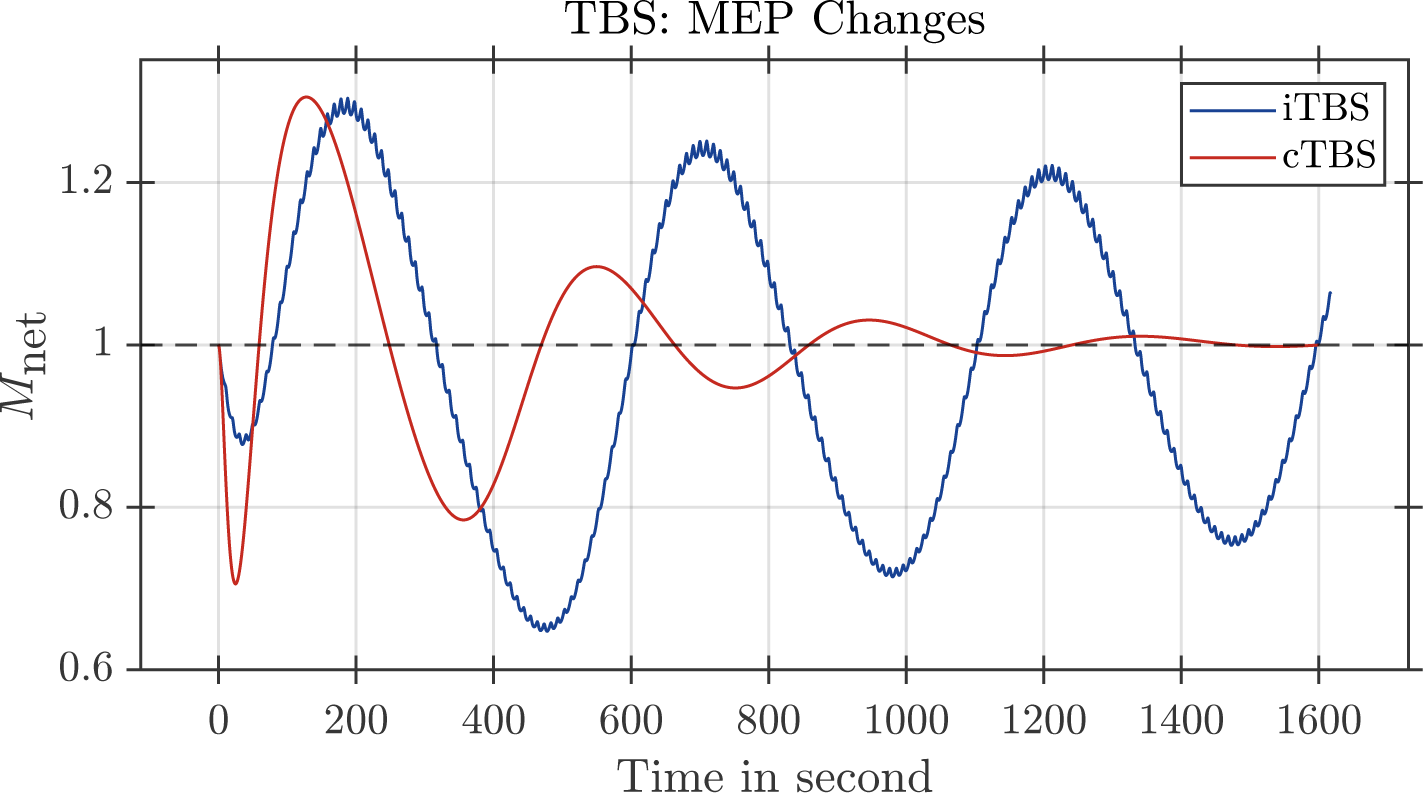
Time evolution of *M*_net_ in responses to continuous application of cTBS and iTBS. A value of *M*_net_ *>* 1 means facilitatory effects on MEP, while *M*_net_ *<* 1 means inhibitory effects on MEP. The blue solid line corresponds to the iTBS protocol, and the solid red line corresponds to the cTBS protocol.

## 5 Discussion

The purpose of this modelling work was to construct a model that can capture the dose-dependency of TBS-induced after-effects and to explore the underlying neural processes involved. Our model is based on the well-established mechanisms of NMDAR-dependent synaptic plasticity and metaplasticity, which are widely accepted in the field. This model is a type of neural mass model that particularly focuses on glutamatergic synapses. The key hypothesis is that the calcium influx through NMDAR simultaneously induces a mixture of facilitatory and inhibitory effects during the magnetic intervention. We constructed a nonlinear model that incorporates these mechanisms and hypotheses. The model was calibrated using experimental data from TBS studies, allowing us to estimate a set of consistent parameters. Although the model successfully described the after-effects induced by prolonged TBS protocols, it is important to note that the model does not provide direct evidence for the underlying mechanisms and prove the proposed mechanisms. Moreover, this model can serve as a theoretical framework for understanding the dose-dependency of TBS-induced after-effects and offers insights into the underlying neural processes. By extrapolating from the calibrated parameters and the observed patterns of TBS-induced after-effects, the model potentially offers a tool for exploring untested protocols and optimising TMS interventions for clinical applications. However, its predictions should be interpreted with caution due to a limited database of TBS protocols.

### Synaptic learning and metaplasticity

This model attributes the changes in MEP size to the bi-directional synaptic induction at glutamatergic synapses in the primary motor cortex. We consider two major forms of bi-directional synaptic plasticity, i.e., LTP and LTD, which can coexist in the same synapse. The key underlying mechanisms for these forms of plasticity involve changes in a-amino-3-hydroxy-5-methyl-4-isoxazole propionic acid receptors (AMPAR) at the postsynaptic sites (Diering and Huganir 2018). Moreover, the involved molecular mechanisms for bi-directional changes in AMPARs differ from each other. Additionally, there are time delays in the single transduction responses between enzymes and their target substances (Baumgärtel and Mansuy 2012; Herring and Nicoll 2016). We simplified the model by representing the two distinct protein cascades involved in LTP and LTD with two exemplary substances, [*Faci*] and [*Inhi*], respectively. This allows us to describe the molecular dynamics activated by the calcium influx through NMDARs during the induction of synaptic plasticity. Furthermore, the dynamics of our model, as described by Equation 3, inherently include a signal latency similar to the biochemical reactions as aforementioned because of the internal properties of a first-order system.

Moreover, metaplasticity is a key mechanism to capture the after-effect reversal induced by prolonged protocols. For example, when *r*_M_ = 0, both iTBS and cTBS will ultimately have a facilitatory effect on MEPs rather than induce oscillations in *M*_net_. In our model, 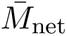 acts as a short-memory device to remember the history of *M*_net_; if *M*_net_ is first facilitated, it becomes easy to induce inhibition for upcoming stimuli and vice versa. This mechanism ensures the stability of MEP learning without diverging to the upper or lower limits of the system. Similarly, the sliding threshold theory proposed by Bienenstock et al. (1982) also introduces extra stability to the Hebbian learning algorithm to avoid unlimited bounds.

### Facilitation and inhibition substances

This model hypothesised that the postsynaptic calcium influx simultaneously triggers the generation of [*Faci*] and [*Inhi*], which is consistent with some findings in the calcium signalling cascades involved in synaptic plasticity. The calcium influx through NMDARs triggers a conformational change in calmodulin (CaM) that can selectively activate both Ca^2+^/calmodulin-dependent protein kinase (CaMKII) and calcineurin (CaN) (Baumgärtel and Mansuy 2012; Herring and Nicoll 2016). CaMKII is a protein kinase that plays a crucial role in LTP, while CaN is a protein serine/threonine phosphatase that is essential for LTD (Baumgärtel and Mansuy 2012; Herring and Nicoll 2016). By modulating the redistribution, trafficking, and phosphorylation/dephosphorylation of AMPARs, both CaMKII and CaN exert control over the bi-directional expression of synaptic plasticity (Baumgärtel and Mansuy 2012; Beattie et al. 2000; Hell 2016; Lee et al. 2000, 1998; Sanderson et al. 2016; Shi et al. 1999). This provides a potential mechanistic link between the calcium influx through NMDARs triggered by stimulus and the generation of [*Faci*] and [*Inhi*] in our model.

CaMKII and CaN can be considered as information decoders that exhibit distinct responses to changes in [Ca^2+^]_i_ post-synaptically (Fujii et al. 2013). The activation of CaMKII is regulated by autophosphorylation at Thr286, and its activity is negatively regulated by dephosphorylation mediated by protein phosphatase 1 (PP1) (Shioda and Fukunaga 2017). Interestingly, the presence of PP1 makes CaMKII highly sensitive to strong calcium signals, and its autophosphorylation exhibits a steep dependence on [Ca^2+^]_i_(Bradshaw et al. 2003). Furthermore, a study by Chang et al. (2017) demonstrated that CaMKII acts as a leaky integrator and repetitive Ca^2+^ elevations significantly activate autophosphorylation of CaMKII over time. This property allows CaMKII to exhibit prolonged activation, improving its ability to integrate Ca^2+^ signals. The nonlinear and ultrasensitive properties of CaMKII highlight its ability to function as a molecular switch that can detect and rapidly respond to subtle changes in [Ca^2+^]_i_ over a narrow range.

Moreover, CaN has a much higher affinity for Ca^2+^/CaM than CaMKII (Klee and Vanaman 1982), and thus the initial elevations in intracellular Ca^2+^ will make CaN-mediated phosphatase activity superior to CaM kinase signalling (Bito et al. 1997). One of these CaN-mediated phosphatases is PP1. Activation of CaN dephosphorylates the inhibitor of PP1, resulting in an increase in PP1 activity (Mulkey et al. 1994). Furthermore, Genoux et al. (2002) demonstrated that PP1 inhibition is associated with increased phosphorylation of CaMKII, thereby affecting synaptic plasticity. However, the inhibition of CaN is not yet fully understood. Several studies have suggested that high Ca^2+^ concentration activates kinases, such as CaMKII and protein kinase C, which can directly phosphorylate and inhibit CaN (Hashimoto and Soderling 1989; Martensen et al. 1989). These findings imply the importance of the kinase-phosphatase crosstalk system in regulating the balance between potentiation and depression during the induction of synaptic plasticity.

In our model, we implemented a Hill function to capture the relationship between [Ca^2+^]_i_ and the growth rate of [*Faci*]. This approach reflects the enhanced ability of [*Faci*] to integrate calcium signals at higher levels of [Ca^2+^]_i_, which is similar to the properties of CaMKII. By using a Hill coefficient of 20, we introduced a steepness factor that ensures that the production rate of [*Faci*] increases rapidly as [Ca^2+^]_i_ exceeds a certain threshold. On the other hand, the growth rate of [*Inhi*] follows an inverse relationship with [Ca^2+^]_i_, which is similar to the properties of CaN mentioned above. It is highest at low [Ca^2+^]_i_ levels and gradually decreases as [Ca^2+^]_i_ increases. This pattern reflects the inhibitory effect of [*Inhi*] on synaptic plasticity when calcium levels are low, and its diminishing influence as calcium levels rise. Reminding of a kinase-phosphatase interaction system, the combined [*Faci*] and [*Inhi*] translate Ca^2+^ signals into either potentiation or depression of synaptic plasticity.

### Further development

The latest more potent neuromodulatory TMS protocols require a large number of parameters, such as pulse strength, pulse/burst frequency, inter-train interval, the number of pulses per burst and trains per session. The parameter selection still remains largely unexplored. This unexplored parameter space suggests a great potential for higher efficacy, speed, and/or effect reliability as well as future optimisation (Caulfield and Brown 2022). This model simplifies modulatory TMS protocols as a series of impulse inputs and thus ignores some important considerations regarding the input variables and their effects on neural populations. One such consideration is pulse strength. Recent evidence has shown that altering the pulse strength may shift the balance between different activated neural populations and therefore affect the after-effects (Lang et al. 2006; Sasaki et al. 2018). However, the calibration of model parameters for the pulse strength is challenging due to the lack of diversity of strength used across studies (Turi et al. 2021).

Another consideration is the waveform of the pulses. Conventional monophasic pulses are known to outcompete biphasic pulses in modulating the primary motor cortex (Arai et al. 2007; Nakamura et al. 2016; Tiksnadi et al. 2020). More advanced and intentionally designed pulse waveforms, such as asymmetric near-rectangular electric fields or monophasic/biphasic pulses, have been found to have different effects on cortical excitability and after-effects. Various studies demonstrated that asymmetric near-rectangular electric field pulses can generate a stronger reduction in excitability and induce longer-lasting aftereffects compared to biphasic pulses (Casula et al. 2018; Goetz et al. 2016; Halawa et al. 2019), while some of these pulses can also have other physical advantages (Ma and Goetz 2023; Zhang et al. 2023). These findings indicate the importance of pulse waveform as a factor in determining the effects of TMS, although the mechanisms explaining how are not known yet and therefore complicated to model. More systematic experimental measurement of neuromodulation and matching with selectivity measurements, such as latency and activation patterns in electroencephalography may inform future models and in turn, allow testable predictions.

Moreover, the signalling pathway from calcium influx to downstream kinase– phosphatase mechanisms is modelled as a still simple nonlinear system in this study. In reality, the signalling cascades involved in synaptic plasticity and metaplasticity are highly complex and involve multiple molecules, pathways, and feedback loops (Chang et al. 2017; Kornijcuk et al. 2020). Gentner et al. (2008) demonstrated that a voluntary muscle contraction before the neuromodulatory intervention can reverse the after-effect induced by cTBS300 without prior muscle contraction, and Wankerl et al. (2010) suggested that L-type voltage-gated calcium channel may be involved mediating after-effect reversal induced by endogenous neuronal activation. Although NMDAR is a major source of calcium influx, the influence of other calcium sources, such as voltage-gated calcium channels and calcium releases from intracellular stores (Grehl et al. 2015), could be added to the model once sufficient experimental data exist to manage the increased number of degrees of freedom such addition would entail. In addition, recent studies have highlighted the potential contributions of GABAergic neurotransmission in the induction of after-effects following neuromodulatory TMS interventions (Di Lazzaro et al. 2002b; Di Lazzaro et al. 2006; Li et al. 2019). This suggests that the interaction between GABAergic and glutamatergic neurons may contribute to the inter-variability observed in TMS neuromodulation experiments. Integrating the interaction between different neurotransmitter systems could provide a more comprehensive modelling structure for the TMS-induced after-effects. Although direct evidence linking TMS to synaptic plasticity mechanisms is still lacking, the observed neurophysiological features of after-effects following TMS appear consistent with those found in synaptic plasticity (Huang et al. 2017). These similarities suggest that TMS may engage similar cellular mechanisms as those involved in synaptic plasticity, albeit with potential additional contributions from other factors.

Furthermore, the model developed in this study is based on the available data and, therefore, its ability to predict the after-effects induced by new TBS protocols may be limited. The majority of studies used cTBS300, cTBS600, and iTBS600 as preferences (Chung et al. 2016; Corp et al. 2020), and only a few studies used TBS protocols over 600 pulses. The model can successfully reproduce the experimental results within the database range, and extrapolating its predictions to protocols beyond the data range should be done with caution. Therefore, future experimental studies and model refinements are necessary to enhance our understanding of the underlying mechanisms and to improve the accuracy and generalisability of the model.

Most importantly, despite the large trend to data sharing in science, also encouraged by funding bodies, most TMS studies still only report rather limited information. MEPs are often provided as linear averages, although it has been intensively studied that MEPs are far from a Gaussian or any other symmetrical distribution so that simple averaging leads to large bias (Goetz et al. 2019, 2022, 2014; Goetz and Peterchev 2012; Nielsen 1996; Roy Choudhury et al. 2011). The limited data availability and the risk of biased statistics also affect the model quality. Accordingly, it appears recommended that future TMS studies improve the statistical analysis and reporting as they limit interpretability and quantitative analysis. In addition to reporting median and further quantiles, full distributions and raw data may substantially contribute to further TMS studies.

## 6 Conclusion

Our modelling approach provides a theoretical framework for understanding the dose-dependency of TBS-induced after-effects and offers insights into the potential underlying neural processes. Although the model has its simplifications and limitations, it quantitatively captures the observed TBS-induced after-effects and brings models one step closer to systematic optimisation of modern neuromodulation protocols, which overwhelm users, for both experimental brain research and clinical applications (Caulfield and Brown 2022). Moreover, this model suggests that the molecular mechanisms of kinases and phosphatases, such as CaMKII and CaN, could be potential candidates for later TMS neuromodulation modelling research. Future studies should aim to incorporate more realistic and complex signalling cascades, consider the involvement of GABAergic neurotransmission, and explore the interaction of different neurotransmitter systems to enhance our understanding of the mechanisms underlying TMS-induced after-effects. In conclusion, this model provides a starting basis for including more complex dynamics at both cellular and molecular levels and potential guidance for flexibly manipulating and optimising TBS parameters with low computational cost, aiding the design of TMS protocols prior to experimental testing.

## Supplementary Material

The list of collected studies is uploaded with this paper. An implementation of the model in MATLAB accompanied by instructions for its use is available at https://github.com/BIOMAKE/Simple_Nonlinear_TBS_Model.

