## supplementary material for "Exploring the Dose-Dependency of After-Effects: A Computational Model for Theta-Burst Transcranial Magnetic Stimulation"

### Database paper list

**Table 1:** Summary of the collected studies for calibration of model parameters.

| Study | Sample Size | Gender ratio | Mean age ± SD (age range) | TBS Protocol | Pulse Strength | Pulse Number | Target Muscle |
|---|---|---|---|---|---|---|---|
| Antal et al. (2010) | 10 | 7 F:3 M | (21 − 32) | iTBS | 80 % AMT | 600 | Right FDI |
| | 5 | 3 F:2 M | (20 − 29) | iTBS | 80 % AMT | 600 | Right FDI |
| Belvisi et al. (2013) | 14 | 3 F:11 M | 41.9 ± 11.36 (23 − 60) | iTBS | 80 % AMT | 600 | Right FDI |
| Brownjohn et al. (2014) | 10 | 1 F:9 M | 26.9 ± 4.7 (22 − 37) | iTBS | 80 % AMT | 600 | Right FDI |
| | 10 | 1 F:9 M | 26.9 ± 4.7 (22 − 37) | cTBS | 80 % AMT | 600 | Right FDI |
| Cheeran et al. (2008) | 9 | 3 F:6 M | 29.3 ± 3 | iTBS | 80 % AMT | 600 | Right FDI |
| | 9 | 3 F:6 M | 28.7 ± 3 | iTBS | 80 % AMT | 600 | Right FDI |
| | 9 | 5 F:4 M | 26.45 ± 5 | cTBS | 80 % AMT | 300 | Right FDI |
| | 9 | 5 F:4 M | 26.45 ± 5 | cTBS | 80 % AMT | 300 | Right FDI |
| Chuang et al. (2014) | 18 | 11 F:7 M | 48.6 ± 12.8 | cTBS | 80 % AMT | 600 | Right FDI |
| Conte et al. (2012) | 15 | - | 68.1 ± 10.2 | iTBS | 80 % AMT | 600 | Right FDI |
| | 7 | - | 65.3 ± 12.1 | cTBS | 80 % AMT | 600 | Right FDI |
| Di Lazzaro et al. (2008) | 12 | - | 63.2 ± 5.3 | iTBS | 80 % AMT | 600 | Right FDI |
| | 12 | - | 63.2 ± 5.3 | cTBS | 80 % AMT | 600 | Right FDI |
| Di Lazzaro et al. (2011) | 10 | - | 26.6 ± 4.1 | iTBS | 80 % AMT | 600 | Left FDI |
| | 10 | - | 26.6 ± 4.1 | cTBS | 80 % AMT | 600 | Left FDI |
| Di Lorenzo et al. (2020) | 12 | - | 71.1 ± 5.9 | iTBS | 80 % AMT | 600 | Right FDI |
| | 12 | - | 71.1 ± 5.9 | cTBS | 80 % AMT | 600 | Right FDI |
| Doeltgen and Ridding (2011) | 14 | 10 F:4 M | 24.5 ± 3.1 | iTBS | 80 % AMT | 600 | Right FDI |
| | 9 | 6 F:3 M | 23.2 ± 3.7 | iTBS | 80 % AMT | 600 | Right FDI |
| | 14 | 10 F:4 M | 24.5 ± 3.1 | cTBS | 80 % AMT | 600 | Right FDI |
| | 9 | 6 F:3 M | 23.2 ± 3.7 | cTBS | 80 % AMT | 600 | Right FDI |
| Doeltgen et al. (2012) | 17 | 10 F:7 M | 23.1 ± 5.1 | cTBS | 80 % AMT | 600 | Right FDI |
| Edwards et al. (2006) | 10 | 3 F:7 M | (26 − 69) | cTBS | 80 % AMT | 300 | Right FDI |
| Fang et al. (2014) | 9 | 4 F:5 M | 24.2 ± 2.0 | cTBS | 80 % AMT | 300 | Right FDI |
| Gamboa et al. (2010) | 14 | 7 F:7 M | (21 − 27) | iTBS | 80 % AMT | 600 | Right FDI |
| | 14 | 7 F:7 M | (21 − 27) | iTBS | 80 % AMT | 1200 | Right FDI |
| | 14 | 7 F:7 M | (21 − 27) | cTBS | 80 % AMT | 600 | Right FDI |
| | 14 | 7 F:7 M | (21 − 27) | cTBS | 80 % AMT | 1200 | Right FDI |
| Gamboa et al. (2011) | 16 | 6 F:10 M | (21 − 27) | iTBS | 80 % AMT | 600 | Right FDI |
| | 16 | 6 F:10 M | (21 − 27) | cTBS | 80 % AMT | 600 | Right FDI |
| Goldsworthy et al. (2012a) | 12 | 6 F:6 M | 23.7 ± 8.1 | cTBS | 80 % AMT | 600 | Right FDI |
| Goldsworthy et al. (2012b) | 12 | 7 F:5 M | 26.3 ± 2.3 | cTBS | 80 % AMT | 600 | Right FDI |
| Guerra et al. (2019) | 18 | 6 F:12 M | 26.1 ± 1.9 | cTBS | 80 % AMT | 600 | Right FDI |
| Hamada et al. (2013) | 56 | 24 F:32 M | 30.3 ± 7.4 (18 − 52) | iTBS | 80 % AMT | 600 | Right FDI |
| | 56 | 24 F:32 M | 30.3 ± 7.4 (18 − 52) | cTBS | 80 % AMT | 600 | Right FDI |
| Hasan et al. (2012) | 9 | 2 F:7 M | 30.3 ± 1.5 | iTBS | 80 % AMT | 600 | Right FDI |
| | 9 | 2 F:7 M | 30.3 ± 1.5 | cTBS | 80 % AMT | 600 | Right FDI |
| He et al. (2021) | 18 | - | - | iTBS | 80 % AMT | 600 | Left FDI |
| Hinder et al. (2014) | 30 | 19 F:11 M | 25.3 ± 8.7 | iTBS | 80 % AMT | 600 | Right FDI |
| Huang et al. (2005) | 9 | - | 33.6 ± 7.8 (23 − 52) | iTBS | 80 % AMT | 600 | Right FDI |
| | 9 | - | 33.6 ± 7.8 (23 − 52) | cTBS | 80 % AMT | 300 | Right FDI |
| | 9 | - | 33.6 ± 7.8 (23 − 52) | cTBS | 80 % AMT | 600 | Right FDI |
| Huang et al. (2007) | 6 | 5 F:1 M | 26 ± 9 | iTBS | 80 % AMT | 600 | Right FDI |
| | 6 | 5 F:1 M | 26 ± 9 | cTBS | 80 % AMT | 300 | Right FDI |

**Table 1:** Summary of the collected studies for calibration of model parameters.

| Study | Sample Size | Gender ratio | Mean age ± SD (age range) | TBS Protocol | Pulse Strength | Pulse Number | Target Muscle |
|---|---|---|---|---|---|---|---|
| Huang et al. (2009) | 8 | 5 F:3 M | $35 \pm 14$ | cTBS | 80 % AMT | 300 | Right FDI |
| Huang et al. (2010a) | 8 | 7 F:1 M | $33.3 \pm 10.3$ | iTBS | 80 % AMT | 600 | Right FDI |
| | 7 | 3 F:4 M | $28.7 \pm 3.6$ | cTBS | 80 % AMT | 600 | Right FDI |
| Huang et al. (2010b) | 9 | 5 F:4 M | $42.7 \pm 12.1$ | cTBS | 80 % AMT | 300 | Right FDI |
| | 9 | 5 F:4 M | $42.7 \pm 12.1$ | cTBS | 80 % AMT | 600 | Right FDI |
| Iezzi et al. (2011) | 10 | 4 F:6 M | $32 \pm 5.03$ | iTBS | 80 % AMT | 600 | Right FDI |
| | 10 | 4 F:6 M | $32 \pm 5.03$ | cTBS | 80 % AMT | 600 | Right FDI |
| Ishikawa et al. (2007) | 10 | 1 F:9 M | $42.3 \pm 6.9$ | cTBS | 80 % AMT | 600 | Right FDI |
| Kimura et al. (2022) | 18 | 5 F:13 M | $21.7 \pm 1.0$ | iTBS | 80 % AMT | 600 | Right FDI |
| Kishore et al. (2012) | 10 | - | $45.6 \pm 7.8$ | iTBS | 80 % AMT | 600 | Right FDI |
| | 10 | - | $45.6 \pm 7.8$ | cTBS | 80 % AMT | 600 | Right FDI |
| Koch et al. (2012) | 14 | - | - | iTBS | 80 % AMT | 600 | Right FDI |
| | 14 | - | - | cTBS | 80 % AMT | 600 | Right FDI |
| Koch et al. (2014) | 10 | 6 F:4 M | $68.3 \pm 5.6$ | iTBS | 80 % AMT | 600 | Right FDI |
| | 10 | 6 F:4 M | $68.3 \pm 5.6$ | cTBS | 80 % AMT | 600 | Right FDI |
| Li Voti et al. (2011) | 21 | - | - | iTBS | 80 % AMT | 600 | Right FDI |
| Mastroeni et al. (2013) | 29 | 29 M | $26.0 \pm 3.2$ | iTBS | 80 % AMT | 600 | Right FDI |
| | 29 | 29 M | $26.0 \pm 3.2$ | cTBS | 80 % AMT | 600 | Right FDI |
| McAllister et al. (2011) | 23 | 13 F:10 M | $27.9 \pm 8.3$ | cTBS | 80 % AMT | 600 | Right FDI |
| McAllister et al. (2013) | 16 | 9 F:7 M | $(19 - 44)$ | cTBS | 80 % AMT | 600 | Right FDI |
| McCalley et al. (2021) | 30 | 20 F:10 M | $24.4 \pm 3.7$ | iTBS | 80 % AMT | 600 | Right APB |
| | 30 | 20 F:10 M | $24.4 \pm 3.7$ | iTBS | 80 % AMT | 1200 | Right APB |
| | 30 | 20 F:10 M | $24.4 \pm 3.7$ | iTBS | 80 % AMT | 1800 | Right APB |
| | 30 | 18 F:12 M | $25.0 \pm 3.4$ | cTBS | 80 % AMT | 600 | Right APB |
| | 30 | 18 F:12 M | $25.0 \pm 3.4$ | cTBS | 80 % AMT | 1200 | Right APB |
| | 30 | 18 F:12 M | $25.0 \pm 3.4$ | cTBS | 80 % AMT | 1800 | Right APB |
| Moliadze et al. (2014) | 12 | - | $25.7 \pm 4.1$ | iTBS | 80 % AMT | 600 | Right FDI |
| Monte-Silva et al. (2011) | 12 | 6 F:6 M | $25.75 \pm 5.11$ | iTBS | 80 % AMT | 600 | Right FDI |
| | 12 | 6 F:6 M | $25.75 \pm 5.11$ | cTBS | 80 % AMT | 600 | Right FDI |
| Mori et al. (2012) | 77 | 46 F:31 M | $38.3 \pm 10.2$ | iTBS | 80 % AMT | 600 | Right FDI |
| | 77 | 46 F:31 M | $38.3 \pm 10.2$ | cTBS | 80 % AMT | 600 | Right FDI |
| Mori et al. (2013) | 13 | 5 F:8 M | $35.5 \pm 9.2$ | iTBS | 80 % AMT | 600 | Right FDI |
| | 13 | 5 F:8 M | $35.5 \pm 9.2$ | cTBS | 80 % AMT | 600 | Right FDI |
| Murakami et al. (2008) | 6 | - | - | iTBS | 80 % AMT | 600 | Right FDI |
| | 6 | - | - | cTBS | 80 % AMT | 600 | Right FDI |
| Oberman et al. (2012) | 20 | 4 F:16 M | $34.9 \pm 16.2$ | iTBS | 80 % AMT | 600 | Right FDI |
| | 20 | 4 F:16 M | $34.9 \pm 16.2$ | cTBS | 80 % AMT | 600 | Right FDI |
| Opie et al. (2013) | 11 | 2 F:9 M | $43.0 \pm 10.3$ | cTBS | 80 % AMT | 600 | Right FDI |
| Orth et al. (2010) | 14 | 9 F:5 M | $(28 - 62)$ | cTBS | 80 % AMT | 300 | Right FDI |
| Pichiorri et al. (2012) | 11 | 3 F:8 M | $31 \pm 8.5$ | iTBS | 80 % AMT | 600 | Right FDI |
| Player et al. (2012) | 16 | 7 F:9 M | - | iTBS | 80 % AMT | 600 | Right FDI |
| Suppa et al. (2008) | 15 | - | $(26 - 45)$ | iTBS | 80 % AMT | 600 | Left FDI |
| | 15 | - | $(26 - 45)$ | cTBS | 80 % AMT | 600 | Left FDI |
| | 5 | - | $(26 - 45)$ | cTBS | 80 % AMT | 600 | Right FDI |
| | 5 | - | $(26 - 45)$ | cTBS | 80 % AMT | 600 | Left FDI |
| Suppa et al. (2011a) | 14 | 3 F:11 M | $60 \pm 11.28$ $(49 - 81)$ | iTBS | 80 % AMT | 600 | Right FDI |
| Suppa et al. (2011b) | 12 | 5 F:7 M | $30 \pm 4.9$ $(25 - 40)$ | iTBS | 80 % AMT | 600 | Right FDI |
| | 12 | 5 F:7 M | $30 \pm 4.9$ $(25 - 40)$ | cTBS | 80 % AMT | 600 | Right FDI |
| Suppa et al. (2014b) | 20 | 10 F:10 M | $56.6 \pm 11.5$ $(36 - 81)$ | iTBS | 80 % AMT | 600 | Right FDI |

**Table 1:** Summary of the collected studies for calibration of model parameters.

| Study | Sample Size | Gender ratio | Mean age ± SD (age range) | TBS Protocol | Pulse Strength | Pulse Number | Target Muscle |
|---|---|---|---|---|---|---|---|
| | 20 | 10 F:10 M | 56.6 ± 11.5 (36 − 81) | cTBS | 80 % AMT | 600 | Right FDI |
| Suppa et al. (2014a) | 20 | 6 F:14 M | 32.8 ± 11.2 | iTBS | 80 % AMT | 600 | Right FDI |
| | 20 | 6 F:14 M | 32.8 ± 11.2 | cTBS | 80 % AMT | 600 | Right FDI |
| Swayne et al. (2009) | 10 | 3 F:7 M | 29.6 ± 4.7 | iTBS | 80 % AMT | 600 | Right FDI |
| Talelli et al. (2007) | 18 | 9 F:9 M | 29.6 ± 3.9 | iTBS | 80 % AMT | 600 | Right FDI |
| | 18 | 9 F:9 M | 29.6 ± 3.9 | cTBS | 80 % AMT | 300 | Right FDI |
| Teo et al. (2007) | 6 | 2 F:4 M | - | iTBS | 80 % AMT | 300 | Right FDI |
| Todd et al. (2009) | 20 | 12 F:8 M | 25 ± 8 | iTBS | 80 % AMT | 600 | Right FDI |
| | 8 | 4 F:4 M | 27 ± 10 | cTBS | 80 % AMT | 600 | Right FDI |
| Vallence et al. (2013) | 18 | 9 F:9 M | 23.3 ± 2.7 | iTBS | 80 % AMT | 600 | Right APB |
| | 18 | 9 F:9 M | 23.3 ± 2.7 | cTBS | 80 % AMT | 600 | Right APB |
| Wu and Gilbert (2012) | 11 | - | - | iTBS | 80 % AMT | 600 | Right FDI |
| Young-Bernier et al. (2014) | 20 | 13 F:7 M | 22.3 ± 3.2 | iTBS | 80 % AMT | 600 | Right FDI |
| | 18 | 9 F:9 M | 70.1 ± 5.6 | iTBS | 80 % AMT | 600 | Right FDI |
| Zafar et al. (2008) | 9 | 5 F:4 M | 21.3 (21 − 26) | iTBS | 80 % AMT | 600 | Right FDI |
| | 9 | 5 F:4 M | 21.3 (21 − 26) | cTBS | 80 % AMT | 600 | Right FDI |
| Zamir et al. (2012) | 10 | 6 F:4 M | 63.1 ± 8.8 (50-75) | iTBS | 80 % AMT | 600 | Right FDI |

**Note:** (a) SD represents standard deviation; (b) iTBS represents intermittent theta-burst stimulation; (c) cTBS is continuous theta-burst stimulation; (d) AMT is active motor threshold; (e) FDI is the first dorsal interosseous muscle; (f) APB is the abductor pollicis brevis muscle.
